# Fluoxetine rescues rotarod motor deficits in *Mecp2* heterozygous mouse model of Rett syndrome via brain serotonin

**DOI:** 10.1101/2020.06.12.147876

**Authors:** Claudia Villani, Giuseppina Sacchetti, Mirjana Carli, Roberto W. Invernizzi

**Affiliations:** Laboratory of Neurochemistry and Behavior, Department of Neuroscience, Istituto di Ricerche Farmacologiche “Mario Negri” IRCCS, Milano, Italy

**Author notes:** **Correspondence** should be addressed to Dr Roberto W. Invernizzi, Istituto di Ricerche Farmacologiche Mario Negri IRCCS, Via Mario Negri 2, 20156 Milano, Italy. **Abbreviations:** 5-hydroxyindoleacetic acid, 5-HIAA; 8-hydroxy-di-n-propylamino tetraline, 8-OH-DPAT; analysis of variance, ANOVA; beam walking, BM; citalopram, CIT; fluoxetine, FLX; heterozygous, Het or Het; insulin growth factor-1, IGF-1; Methyl-CpG-binding protein 2, Mecp2; p-chlorophenylalanine, pCPA; Rett syndrome, RTT; rotarod RR; Selective serotonin reuptake inhibitors, SSRIs; serotonin, 5-HT; tryptophan hydroxylase-2, Tph2; vehicle, Veh; wild type, WT.

**Keywords:** Fluoxetine, *Mecp2*, Motor dysfunction, Rett syndrome, Serotonin

## Abstract

Motor skill is a specific area of disability of Rett syndrome (RTT), a rare disorder occurring almost exclusively in girls, caused by loss-of-function mutations of the X-linked *methyl-CpG-binding protein2* (*MECP2*) gene, encoding the MECP2 protein, a member of the methyl-CpG-binding domain nuclear proteins family. Brain 5-HT, which is defective in RTT patients and *Mecp2* mutant mice, regulates motor circuits and SSRIs enhance motor skill learning and plasticity.

In the present study, we used heterozygous (Het) *Mecp2* female and *Mecp2*-null male mice to investigate whether fluoxetine, a SSRI with pleiotropic effects on neuronal circuits, rescues motor coordination deficits. Repeated administration of 10 mg/kg fluoxetine fully rescued rotarod deficit in *Mecp2* Het mice regardless of age, route of administration or pre-training to rotarod. The motor improvement was confirmed in the beam walking test while no effect was observed in the hanging-wire test, suggesting a preferential action of fluoxetine on motor coordination. Citalopram mimicked the effects of fluoxetine, while the inhibition of 5-HT synthesis abolished the fluoxetine-induced improvement of motor coordination. *Mecp2* null mice, which responded poorly to fluoxetine in the rotarod, showed reduced 5-HT synthesis in the prefrontal cortex, hippocampus and striatum, and reduced efficacy of fluoxetine in raising extracellular 5-HT as compared to female mutants. No sex differences were observed in the ability of fluoxetine to desensitize 5-HT_1A_ autoreceptors upon repeated administration. These findings indicate that fluoxetine rescues motor coordination in *Mecp2* Het mice through its ability to enhance brain 5-HT and suggest that drugs enhancing 5-HT neurotransmission may have beneficial effects on motor symptoms of RTT.

## 1. Introduction

Mutations of the methyl-CpG-binding protein2 (*Mecp2*), an X-linked gene, is the leading cause of Rett syndrome (RTT), a rare disease affecting females from the first/second year of life (Amir et al., 1999). Affected patients develop a variety of symptoms including motor and cognitive deficits, autistic-like features, convulsions and breathing irregularities (Chahrour and Zoghbi, 2007; Hagberg et al., 1983). Motor deficits, such as ataxia, apraxia and loss of motor skill are prominent in RTT and are consistently reproduced in mice with *Mecp2* deletion or truncation regardless of the sex and background strain (Lombardi et al., 2015). Although hypotrophic fibers, disorganized morphology and altered insulin growth factor-1 signaling is found in muscles of *Mecp2*-null mice, specific deletion of *Mecp2* in skeletal muscles does not reproduce these alterations suggesting that motor phenotype in RTT mice reflects non cell-autonomous mechanisms (Conti et al., 2015). Consistently, conditional deletion of *Mecp2* in nervous tissue and in specific neuronal types of the mouse brain reproduced motor skill deficits typically observed in male *Mecp2*-null and heterozygous (Het) female mice (Chao et al., 2010; Ito-Ishida et al., 2015; Meng et al., 2016; Ure et al., 2016). Therapeutic targets and potential pharmacological strategies are currently being explored, but at present no cure or disease-modifying therapy is available for RTT (Katz et al., 2012). Studies aimed at restoring *Mecp2* function in RTT adult mice through gene therapy, achieved nearly complete reversal of most symptoms including motor abnormalities in *Mecp2*-null and *Mecp2* Het mice (Garg et al., 2013; Guy et al., 2007; Robinson et al., 2012) suggesting that therapeutic intervention for RTT patients can be effective even after symptoms onset. However, most of the pre-clinical studies testing potential pharmacological treatments for RTT did not achieved the complete recovery of motor function (Braun et al., 2012; Buchovecky et al., 2013; Krishnan et al., 2015; Mellios et al., 2014; Nag and Berger-Sweeney, 2007). Thus, the identification of better drugs for RTT is needed.

Low levels of brain serotonin (5-HT), tryptophan hydroxylase-2 (Tph2), the rate-limiting enzyme in brain 5-HT synthesis, and cerebrospinal fluid 5-hydroxyindoleacetic acid (5-HIAA), the main 5-HT metabolite, are found in RTT patients (Riederer et al., 1985; Samaco et al., 2009; Zoghbi et al., 1989). Likewise, reduced levels of 5-HT and 5-HIAA are found in several brain regions of male *Mecp2*-null mice (Ide et al., 2005; Studies in conditional knock-out mice to remove *Mecp2* gene specifically from serotonergic neurons indicate that the loss of *Mecp2* causes a cell-autonomous reduction of brain 5-HT levels and *Tph2* expression (Samaco et al., 2009).

5-HT regulates motor networks facilitating motor skill, motor cortex plasticity and motor output (Vitrac and Benoit-Marand, 2017). Selective 5-HT reuptake inhibitors (SSRIs) enhance motor skill learning and plasticity (Batsikadze et al., 2013; Gerdelat-Mas et al., 2005; Loubinoux et al., 2005), which are defective in RTT patients and *Mecp2* mutant mice (De Filippis et al., 2015; FitzGerald et al., 1990). Increasing brain 5-HT with citalopram improves the CO_2_ chemosensitivity in *Mecp2*-null mice (Toward et al., 2013). In addition, fluoxetine enhances the levels of MeCP2 protein in several brain regions of the rat and in the hippocampus of Ts65Dn mice, a mouse model of Down syndrome (Cassel et al., 2006; Stagni et al., 2015). Thus, in the present study we set out to assess the efficacy of fluoxetine, a SSRI currently used in the therapy of depression, to rescue motor impairment, a specific area of disability of RTT, which can be reliably modeled in mice regardless of sex, age and type of mutation (Lombardi et al., 2015). The role of 5-HT in the motor effects of fluoxetine was assessed by investigating whether a) brain 5-HT depletion by p-chlorophenylalanine (pCPA) affected fluoxetine-induced motor improvement, b) citalopram, another SSRI structurally not related to and by far more selective than fluoxetine (Popik, 1999), mimicked the motor-improving effects of fluoxetine, c) sex-dependent alteration of 5-HT synthesis, release and auto-inhibitory feed-back mechanism occurs under basal conditions and in response to acute and chronic administration of fluoxetine

## 2. Materials and Methods

### 2.1 Experimental design and statistical analysis

Test cohorts composed of *Mecp2* Het (n= 209), *Mecp2*-null mice (n=68) and respective WT (151 females and 64 males) littermate controls were used to assess the ability of fluoxetine and citalopram to counteract motor deficits caused by *Mecp2* deletion using the accelerating rotarod, beam walking and hanging-wire tests, and the role of 5-HT in fluoxetine action. The number of mice/group was established according to recommended guidelines (Katz et al., 2012; Kilkenny et al., 2010), and previous assessments in *Mecp2*-null mice (Villani et al., 2016). Mice were allocated to experimental groups by simple randomization using a freely available software (Statpages.org). Experimenters blind to treatment assignment, assessed motor performance in various behavioral tests.

The primary end-point of the study was to establish whether fluoxetine rescued motor deficit in the rotarod. All statistical analyses were performed by GraphPad Prism version 7.02 for Windows (GraphPad Software, CA) or StatView 5.0.1 (SAS Institute, NC) and all data were expressed as mean ± SEM. For rotarod data, statistical analysis was applied to the mean latency to fall on four daily trials. Two-way ANOVA, followed by Sidak’s test was used to assess the effect of genotype, treatment and their interaction on the latency to fall from rotarod and the number of slips in the beam walking test. Repeated measures ANOVA, with genotype and treatment as between-subject factors and days on treatment as within-subject factor, followed by two-way ANOVA at single time points was used to analyze the onset of fluoxetine effect on the rotarod. The effects of training on the rotarod in naïve *Mecp2* Het and WT mice, the effects of fluoxetine on extracellular 5-HT and 8-OH-DPAT-induced hypothermia was also analyzed by repeated measures ANOVA with genotype or sex and treatment as between-subject factors and time as within-subject factor. Gaussian distribution was formally tested only for rotarod data, the primary end-point of the study, with the D’Agostino-Pearson normality test included in the Graph Pad Prism 7.02 package (D’Agostino, 1986). The analysis confirmed the normal distribution of the latency to fall data with the exception of rotarod data obtained in mice given 20 mg/kg fluoxetine in which n<8 precluded the application of the test. A small departure from normality was observed for the group of mice treated with vehicle + pcpa (p=0.0304). The scores obtained in the hanging-wire test were compared by Dunn’s test for comparison between single means. Differences in 5-HTP accumulation between WT and mutant mice were assessed by Student’s t-test.

### 2.2 Mice breeding and husbandry

*Mecp2* Het female (*Mecp2*^−/+^) and WT male (*Mecp2*^+/y^) mice on C57BL/6J background were purchased from The Jackson Laboratory (Bar Harbor, ME). As *Mecp2*-null mice do not mate, breeding pairs were composed by female *Mecp2* Het and C57BL/6J male mice from the colony, in a 2:1 ratio, to generate all the genotypes needed in the experiments. Mice were maintained in a specific pathogen free animal facility under a 12h light/dark cycle (light on at 7:00 a.m.) at 21 ± 2°C and 55 ± 5% relative humidity, with food (Teklad Global 2018S; Envigo, Udine, Italy) and filtered tap water freely available. At weaning, mice were genotyped using standard procedures, separated and group-housed according to sex and provided with environmental enrichment consisting of a colored plastic shelter in each home cage. Nesting material (wood-wool) was provided to breeding cages.

### 2.3 Ethical statement

The *“Istituto di Ricerche Farmacologiche Mario Negri IRCCS”* adheres to the principle set out in the following law, regulations, and policies governing the care and use of laboratory animals: Italian Governing Law (D.lgs.26/2014; Authorization n. 19/2008-A issued March 6, 2008 by Ministry of Health); “Mario Negri” Institutional Regulations and Policies providing internal authorization for persons conducting animal experiments (Quality Management System Certificate – UNI EN ISO 9001:2008 – Reg. N° 6121); the NIH Guide for the Care and Use of Laboratory Animals (2011 edition) and EU directives and guidelines (EEC Council Directive 2010/63/UE).

### 2.4 Rotarod

Mice were pre-trained on the rotarod (Ugo Basile, Gemonio, Italy) at 5 weeks of age (four daily trials for 3 consecutive days; 4-40 rpm in 300 s). At the end of each trial, mice were returned to the home cage for at least 15 min resting before the next trial. The same procedure was used in the test sessions. A cohort of pre-trained mice (21 WT and 24 *Mecp2* Het), was used to assess the effect of 14-day treatment with fluoxetine. A second cohort of pre-trained *Mecp2* Het (n=32) and WT (n= 21) mice was repeatedly exposed to daily rotarod session 2h and 24h and 3, 7 and 14 days after treatment. These mice were also assessed in the hanging-wire test. Another cohort of *Mecp2* Het mice, repeatedly injected with fluoxetine or water for 17 days, received pCPA from day 15 to 17 and were tested in the rotarod 24 h after the last dose of pCPA (44 *Mecp2* Het). Twenty-one Het and 21 WT female mice were treated with citalopram in the drinking water or plain water and assessed in the rotarod and beam walking test after 14 and 15 days of treatment, respectively.

Because of the short life span and early onset of the motor deficits, male *Mecp2* null mice (n=37) and WT (n=37) were not pre-trained on the rotarod but tested immediately before the beginning of fluoxetine treatment and after 14 days of treatment. Except if otherwise specified, all mice were tested on the rotarod about 24 h after the last dose of the chronic schedule.

Three separate cohort of mice were used to investigate sex-related difference in 5-HT synthesis (7 WT and 6 *Mecp2*-null male mice and 6 WT and 6 *Mecp2*-het female mice); the effects of fluoxetine on extracellular 5-HT (9 WT and 10 *Mecp2*-null male mice and 9 WT and 11 *Mecp2*-het female mice) and the hypothermic response to 8-OH-DPAT (12 WT and 13 *Mecp2*-null male mice and 14 WT and 11 *Mecp2*-het female mice).

### 2.5 Beam walking

A wood plank with a flat surface 8 mm wide and 80 cm long, resting on 2 poles (50 cm above the ground) was used. Training consisted in two consecutive sessions during which mice were positioned at one end of the beam and allowed to cross the beam length, with gentle guiding or prodding as needed, until they cross readily. On the test phase, the number of slips is recorded on 3 consecutive trials, with an intertrial interval of at least 15 min.

### 2.6 Hanging-wire

A stainless steel wire 2 mm thick and 640 mm long was suspended horizontally to two vertical stands at 30 cm height above the ground. A mouse was handled by the tail, suspended by the forelimbs on the middle of the wire and the timer started. When the animal reached one of the platforms positioned at the ends of the wire the timer was stopped and a second trial was started by hanging the mouse on the middle of the wire. If the animal fell, the timer was stopped, the initial score (ten) was diminished by 1 and the procedure was restarted. The test lasted for 180 s or for a maximum of 10 falls.

### 2.7 Drug treatment

Fluoxetine (Casen, Recordati SL, Spain) was purchased from a local pharmacy, dissolved in pyrogen-free water (10 mL/kg) and injected intraperitoneally (i.p.) at 10 or 20 mg/kg (as salt), between 8:00 and 10:00 AM, during the light phase of the light-dark cycle, starting on postnatal day (P) 35 in males and on P63 or P77 in females. Two different groups of mice were given fluoxetine (10 mg/kg/day) or citalopram (20 mg/kg/day) in drinking water. Mean liquid intake was similar in mice receiving fluoxetine, citalopram or plain water (3.6±0.2, 4.1±0.1 and 3.9±0.2 mL/day, respectively). This information and the measurement of average body weight for each genotype, were used to determine the drug concentration required to provide the established daily doses. Assessment of the stability of drug solutions showed no changes over 3 weeks storage at room temperature in drinking bottles (data not shown). pCPA ethylester hydrochloride (100 mg/kg of the free base; Sigma-Aldrich, Italy) was dissolved in water and administered by gavage from day 15 to 17 of repeated fluoxetine treatment, with the last dose of both drugs given 24 h prior to behavioral testing. A timeline of treatment schedules and behavioral assessment is provided in each figure.

### 2.8 5-HTP assays

Brain samples were homogenized in 10 volumes of 0.1 M HClO_4_ containing 0.1% Na_2_EDTA and 0.05% Na_2_S_2_O_5_, stored in minced ice for 30 min and centrifuged at 10000 rpm for 10 min with a Sigma 1-13 centrifuge (Celbio, Italy). Supernatant was filtered through 0.45 μm syringe filter (Perkin Elmer, Italy) and injected into the HPLC coupled to electrochemical detection. 5-HTP was separated by a reverse phase column (Supelcosil LC18-DB 3 µm, 75 × 3 mm, Supelchem, Italy) with a mobile phase consisting of 88 mM NaH_2_PO_4_ monohydrate (VWR, Italy); 0.4 mM sodium octan sulfonate (Sigma, Italy); 0.3 mM Na_2_EDTA (Sigma, Italy). The pH of the solution was adjusted to 2.8 with 85% M H_3_PO_4_ after the addiction of 7% CH_3_OH. The mobile phase was pumped with a LC-20AD (Shimadzu, Italy) isocratic pump at 0.5 mL/min. The electrochemical detector (Coulochem II; ESA, MA, USA) was equipped with a 5011 analytical cell (E_1_= 0 mV; E_2_= +350 mV).

### 2.9 Intracerebral microdialysis

Concentric dialysis probes were made with Cuprophan membrane (216-μm outer diameter, 3,000 Da cutoff, Sorin Biomedica, Italy), essentially as described elsewhere (Robinson and Whishaw, 1988). The membrane exposed to the brain tissue was 3 mm long. Female mice were anesthetized with ketamine/medetomidine 75/1 (mg/kg i. p.) or 3% isoflurane, while most male mice were anesthetized with 3% isoflurane for a rapid recovery post-anesthesia. Mice were positioned on a stereotaxic frame (model 900; Kopf Instruments, USA) and the dialysis probe was lowered into the mPFC and secured to the skull with anchorage screws and dental cement. The stereotaxic coordinates referred to the tip of the probe were: AP +3.7 and L ±0.7 mm from bregma and DV −4.8 mm from dura surface (Franklin and Paxinos, 1997). About 20 h after surgery, the probes were perfused with artificial cerebrospinal fluid (composition in mM: NaCl 140, CaCl_2_ 1.26, KCl 3, MgCl_2_ 1, Na_2_HPO_4_ 1.2, glucose 7.2; pH 7.4 with 0.6 M NaH_2_PO_4_) at 1 μl/min with a CMA/100 pump (CMA Microdialysis AB, Sweden). Once the extracellular 5-HT concentrations were stable (at least three consecutive samples differing less than 20% from the mean basal value), mice were given a challenge dose of fluoxetine (10 mg/kg i.p.). Samples of dialysate were collected every 20 min in autosampler vials containing 2 μL of an antioxidant mixture (composition: acetic acid 0.1 M, Na_2_ EDTA 0.27 mM, L-cysteine 3.3 mM, ascorbic acid 0.5 mM; dilution 1:50) and stored at 5°C until analysis. 5-HT in the dialysate was measured by HPLC coupled to electrochemical detection, (Invernizzi, 2013).

### 2.10 8-OH-DPAT-induced hypothermia

8-OH DPAT is a selective 5-HT_1A_ receptor agonist causing hypothermia mediated by the stimulation of 5-HT_1A_ autoreceptors (Goodwin et al., 1985). To assess the sensitivity of 5-HT_1A_ auto-receptors after repeated treatment with fluoxetine, *Mecp2*-null, *Mecp2* Het and WT mice received 0.2 mg/kg 8-OH DPAT subcutaneously (s.c). This dose was chosen on the basis of a dose-response curve (data not shown) because it causes an intermediate reduction of body temperature that allow to detect either potentiation or reduction of 8-OH-DPAT effects. Core temperature was measured by a BAT-12 Microprobe Thermometer equipped with a RET-3 rectal probe for mice (Physitemp, USA), every 15 minutes over a 2 h period. Temperature measurements were taken between 8:00 and 12:00 A.M.

## 3. Results

### 3.1 Effect of pre-training in the rotarod and age-dependent deficits in the rotarod and hanging-wire tests in Mecp2 Het and WT mice

Five weeks old *Mecp2* Het and WT mice rapidly learned the accelerated rotarod task (Fig. 1A). Repeated measures ANOVA applied to the mean latency to fall of the 4 daily trials (Fig. 1B) showed significant effect of days (F (2,126) = 11.82, p < 0.0001), but not genotype (F(1,126) = 3.483, p = 0.0643) and interaction between days and genotype (F(2,126) = 0.3606, p = 0.698) indicating that the ability to learn the task was not significantly affected by *Mecp2* gene deletion in young *Mecp2* Het mice. Post-hoc analysis showed a significant increase in the latency to fall between day 1 and 2 and day 1 and 3 of training, while no significant differences were observed between day 2 and 3 (Fig. 1B).

**Fig. 1.**
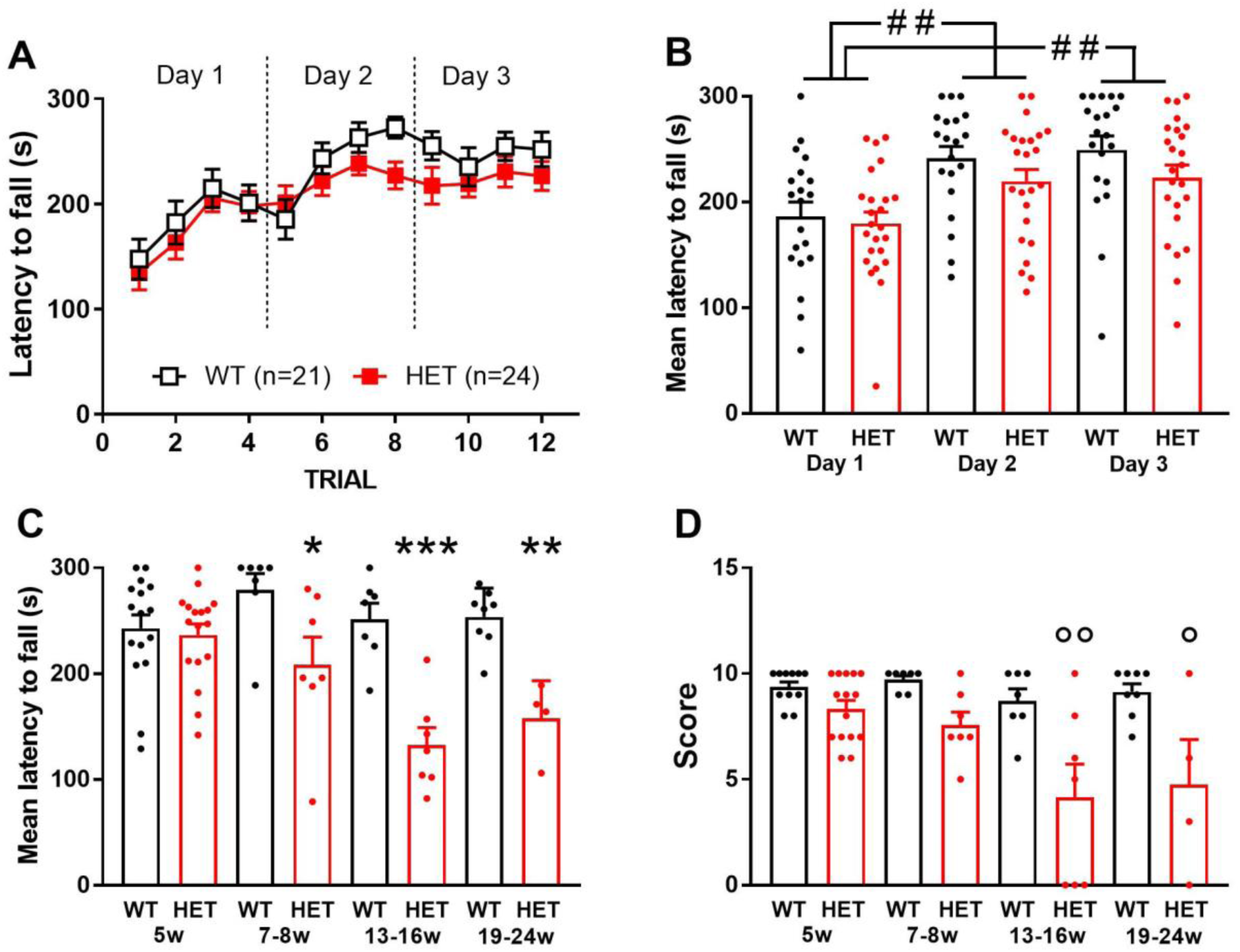
Effect of repeated training on the rotarod and age-dependent performance of untreated WT and Mecp2 Het mice in the rotarod and hanging-wire tests. Panel A describes the trial by trial improvement in latency to fall over 3 consecutive days of training on the rotarod, in 5 weeks old WT and *Mecp2* Het (HET) female mice. Analysis of the mean latency to fall (panel B) show a significant improvement in performance between day 1 and 2, day 1 and 3, but no differences between day 2 and 3 in both genotypes. Panels C and D show the age-dependent performance of WT and *Mecp2* Het mice in the rotarod and hanging-wire test, respectively. Different group of mice were used for each age. Data in panels C and D are the mean ± SEM of 3 consecutive days of trials (4 trials/day). Black and red dots and histograms in panels B, C and D represent individual data points and the mean performance of the group, respectively. The number of mice per group is shown in parentheses. #p = 0.0005; ##p < 0.0001 vs. day 1 (Sidak’s test). *p = 0.0222, **p = 0.0045, ***p < 0.0001 vs. WT; °p = 0.0034, °°p = 0.0004 vs. WT (Sidak’s test).

Using different groups of *Mecp2* Het and age-matched WT mice for each age range, we observed an age-dependent worsening in the performance of rotarod (Fig. 1C) and hanging-wire (Fig. 1D) tests. Rotarod performance (mean latency to fall over 3 days of training) was already reduced in *Mecp2* Het mice aged 7-8 weeks and further reduced in 13-16 and 19-24 weeks old *Mecp2* Het mice. Two-way ANOVA showed highly significant effects of age (F(3,65)= 4.928; p = 0.0038), genotype (F(1,65)= 37.75; p < 0.0001) and interaction between age and genotype (F(3,65)= 6.193; p = 0.0009). Hanging-wire performance was impaired in *Mecp2* Het mice aged 13-16 and 19-24 weeks, whereas no significant differences were observed between *Mecp2* Het and WT mice aged 7-8 weeks. ANOVA showed significant effects of age (F(3,58)= 5.709; p = 0.0017), genotype (F(1,58)= 32.3; p < 0.0001) and interaction between age and genotype (F(3,58)= 3.105; p = 0.0334).

### 3.2 Effect of repeated administration of fluoxetine and citalopram on motor coordination in Mecp2 Het mice

At the end of the 2-week treatment, a complete rescue of the motor deficit was observed in *Mecp2* Het mice receiving 10 mg/kg fluoxetine daily (Figs. 2A and 2B).

**Fig. 2.**
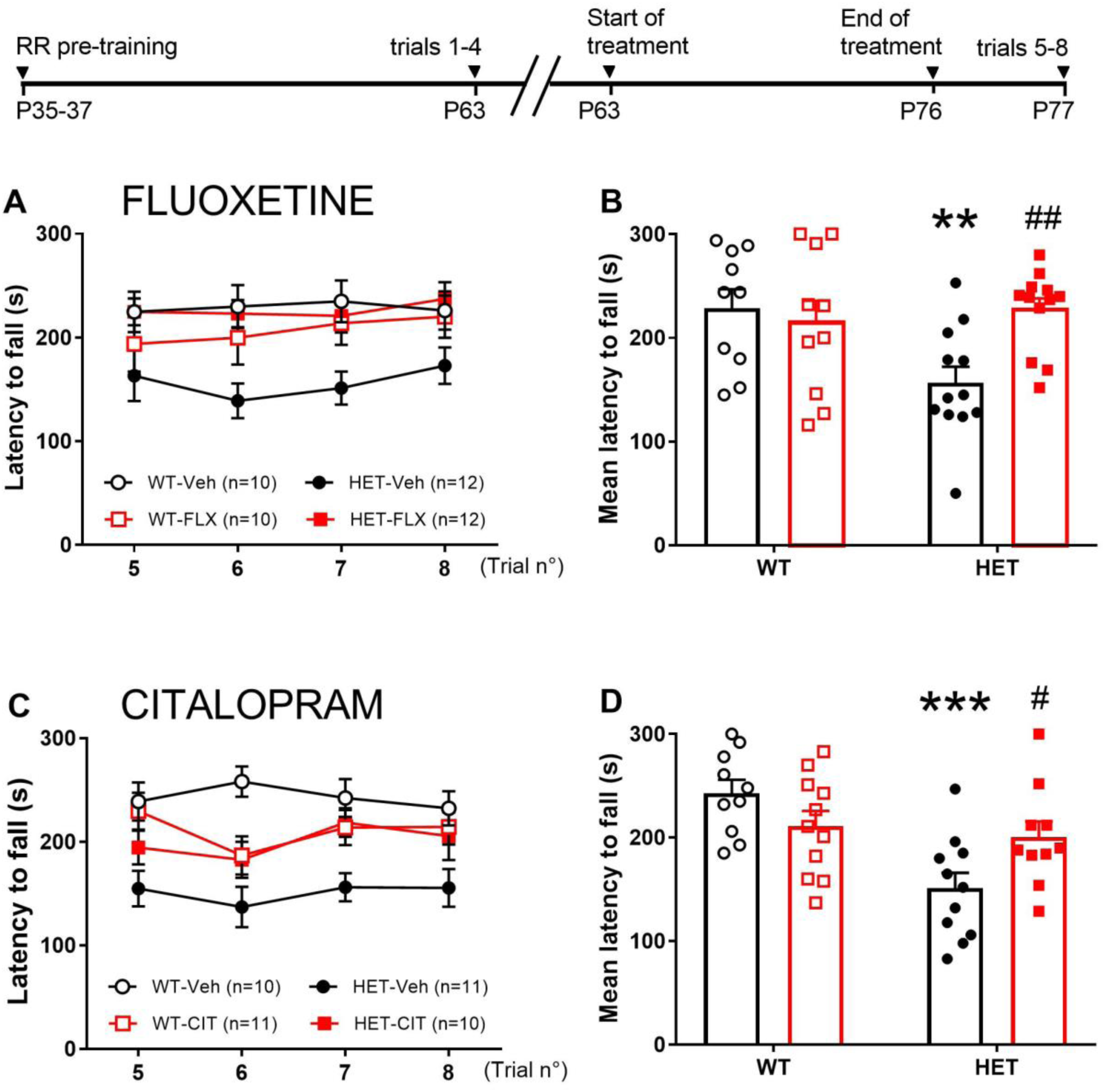
Effect of 14-day treatment with fluoxetine and citalopram on rotarod performance in Mecp2 Het mice. Upper panels show the trial by trial (panel A) and the mean latency to fall (panel B) in *Mecp2* Het and WT mice treated with vehicle (Veh) or fluoxetine. The same parameters measured in mice receiving citalopram (CIT) in the drinking water are shown in panels C and D. The horizontal bar on top of the figure shows the schedule of treatment and the timing of behavioral assessment. Data are mean ± SEM. Symbols and histograms in panels B and D represent individual data points and the mean of the group, respectively. The number of mice per group is shown in parentheses. *p = 0.008 and **p < 0.0001 vs. respective WT-vehicle; #p = 0.0413, ##p = 0.0069 vs. HET-Veh (Sidak’s test). RR, rotarod testing; P, postnatal day

Two-way ANOVA showed a significant interaction between treatment and genotype (F(1,40) = 6.466, p = 0.015), but not genotype (F(1,40) = 3.164, p = 0.083) and treatment (F(1,40) = 2.726, p = 0.107). Post-hoc analysis confirmed that *Mecp2* Het mice treated with fluoxetine remained significantly longer on the rotarod than those receiving vehicle, reaching a performance similar to that observed in WT mice. Fluoxetine had no significant effects on the latency to fall in WT mice.

Two-week administration of 20 mg/kg/day citalopram in the drinking water significantly improved rotarod performance in *Mecp2* Het mice (Figs. 2C and 2D). Two-way ANOVA showed a significant interaction between treatment and genotype (F(1,38) = 7.864, p = 0.0079), genotype (F(1,38) = 12.6, p = 0.001) but not treatment (F(1,38) = 0.3655, p = 0.549).

The number of slips in the beam walking test was higher in *Mecp2* Het than in WT mice and fluoxetine significantly reduced them (Fig. 3A; treatment F(1,32) = 5.039, p = 0.0318; genotype F(1,32) = 39.74, p < 0.00001; and interaction between genotype and treatment (F(1,32) = 5.31, p = 0.0278). Likewise, citalopram reduced the number of slips in *Mecp2* Het mice (Fig. 3B). ANOVA showed a significant effect of treatment (F (1,38) = 8.196, p = 0.0068), genotype (F (1,38) = 21.42, p < 0.0001) and their interaction (F (1,38) = 6.37, p = 0.0159).

**Fig. 3.**
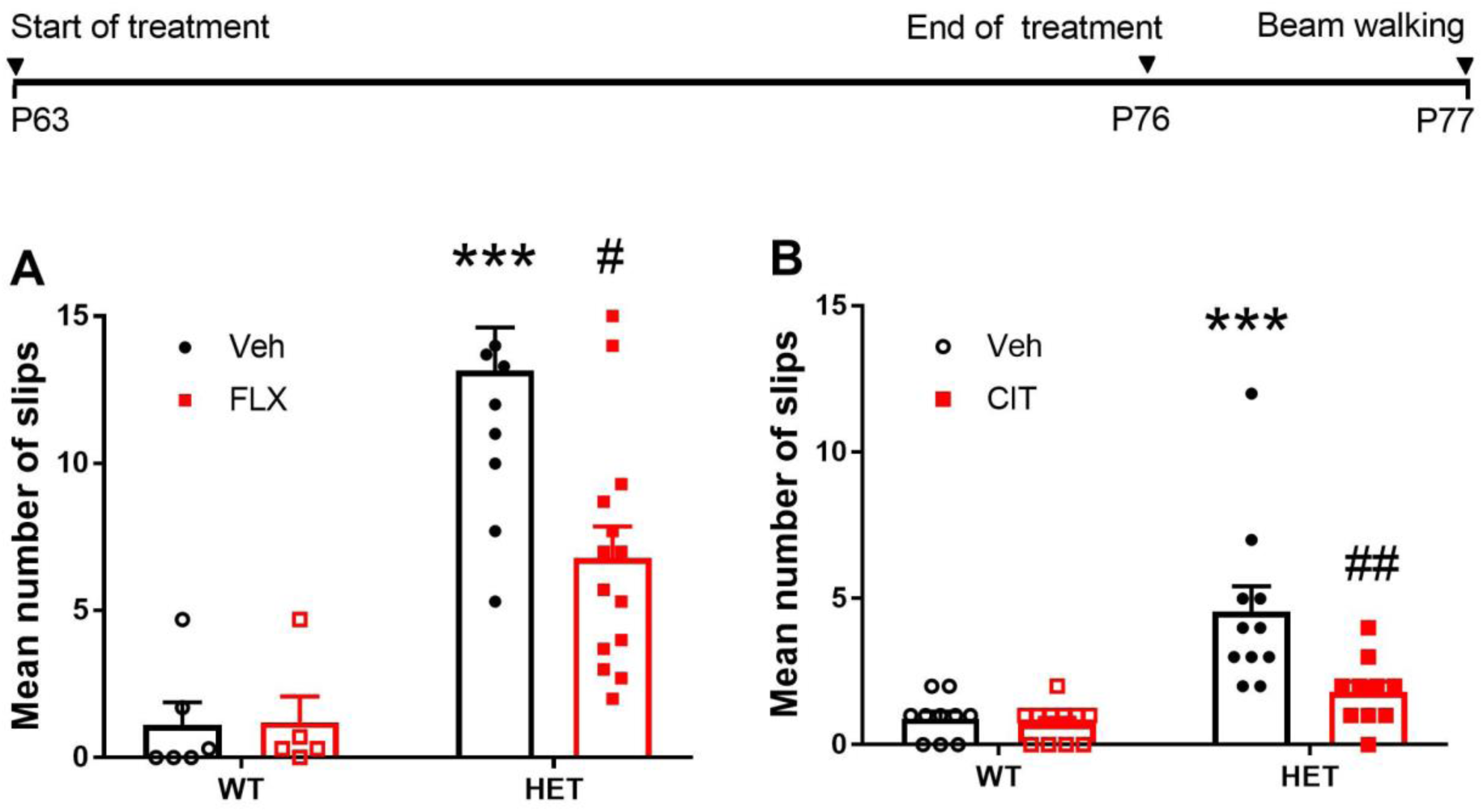
Effect of fluoxetine and citalopram on beam walking performance in Mecp2 Het and WT mice. Panels A and B show the effect of fluoxetine and citalopram, respectively. The horizontal bar on top of the figure illustrate the schedule of treatment and the timing of behavioral assessment. Black and red symbols in panels A and B represent individual data points. Histograms are the mean number of slips over 3 consecutive trials ± SEM of 10-11 mice per group. ***p < 0.0001 vs. WT-H_2_O; #p = 0.0016, ##p = 0.003 vs. HET-Veh (Sidak’s test). BW, beam walking; P, postnatal day.

*Mecp2* Het mice showed a reduced score in the hanging-wire test as compared to WT mice (WT/Vehicle 8.6 ± 0.7, n=11 *Mecp2* Het/vehicle 4.9 ± 0.8, n=16; p < 0.0001, Dunn’s test). Two-week treatment with fluoxetine did not significantly improve the ability of *Mecp2* Het mice to perform the test (WT/fluoxetine 9.1 ± 0.5, n=10, *Mecp2* Het/fluoxetine 5.3 ± 0.8, n=16; p < 0.01, Dunn’s test; data not shown).

Fig. 4 shows the onset of fluoxetine effect on the rotarod and 3 and 7 days after drug suspension. ANOVA showed a highly significant interaction between genotype, treatment and days on treatment (F(4,196) =4.786, p = 0.0011). Further analysis at each time point showed that fluoxetine prolonged the latency to fall in *Mecp2* Het mice after 7 (F(1,49) = 8.002, p = 0.0068) and 14 (F(1,49) = 6.438, p = 0.0144) days but had no effect after 2h, 24 h and 3 days of treatment in *Mecp2* Het mice and at any times after administration in WT mice. In *Mecp2* Het mice, the effect of fluoxetine disappeared 3 days after drug suspension.

**Fig. 4.**
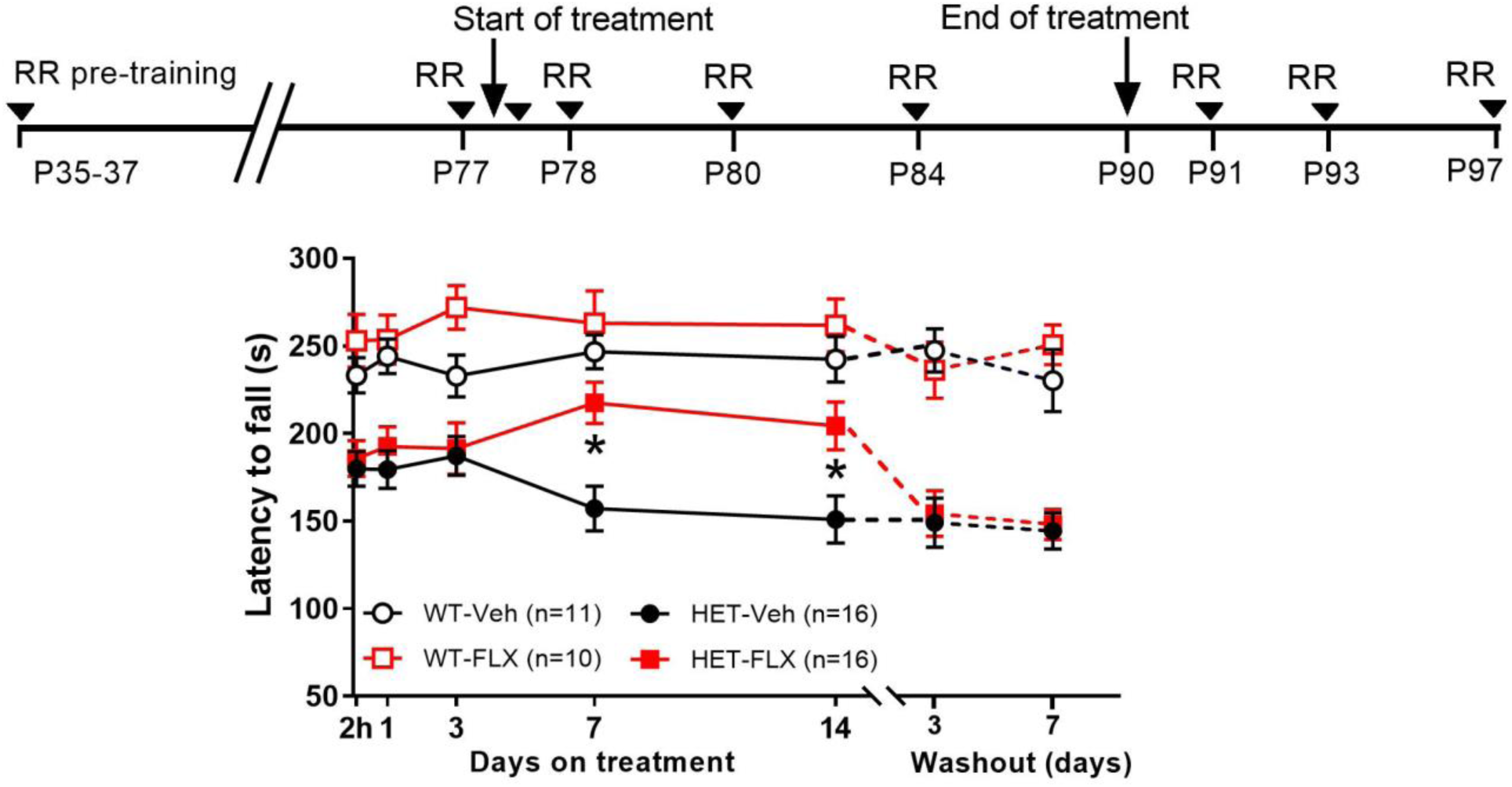
Onset and duration of fluoxetine effect in the rotarod in Mecp2 Het and wild type mice. Mice received 10 mg/kg IP fluoxetine (FLX) or vehicle (H_2_O) daily and were repeatedly exposed to rotarod during treatment and after treatment suspension as indicated by arrowheads in the timeline above the graph. Except for the first time point, mice were exposed to rotarod 24 h after the last dose. Each time-point is the mean ± SEM of 4 daily trials. The number of mice per group is shown in parentheses. Asterisks indicate p < 0.05 vs. HET-Veh (Sidak’s test). RR, rotarod testing; P, postnatal day

### 3.3 Inhibition of 5-HT synthesis prevents fluoxetine ability to rescue rotarod deficit

*Mecp2* Het mice receiving 10 mg/kg fluoxetine for 14 days showed an improved response in the rotarod as compared to those receiving vehicle and pCPA abolished the effect of fluoxetine (Fig. 5). ANOVA showed a significant effect of fluoxetine (F(1,40) = 13.98, p = 0.0006), pCPA (F(1,40) = 7.037, p = 0.0114) and interaction between fluoxetine and pCPA (F(1,40) = 8.103, p = 0.0069). pCPA had no significant effect on rotarod performance in *Mecp2* Het mice treated with vehicle and in WT mice (data not shown).

**Fig. 5.**
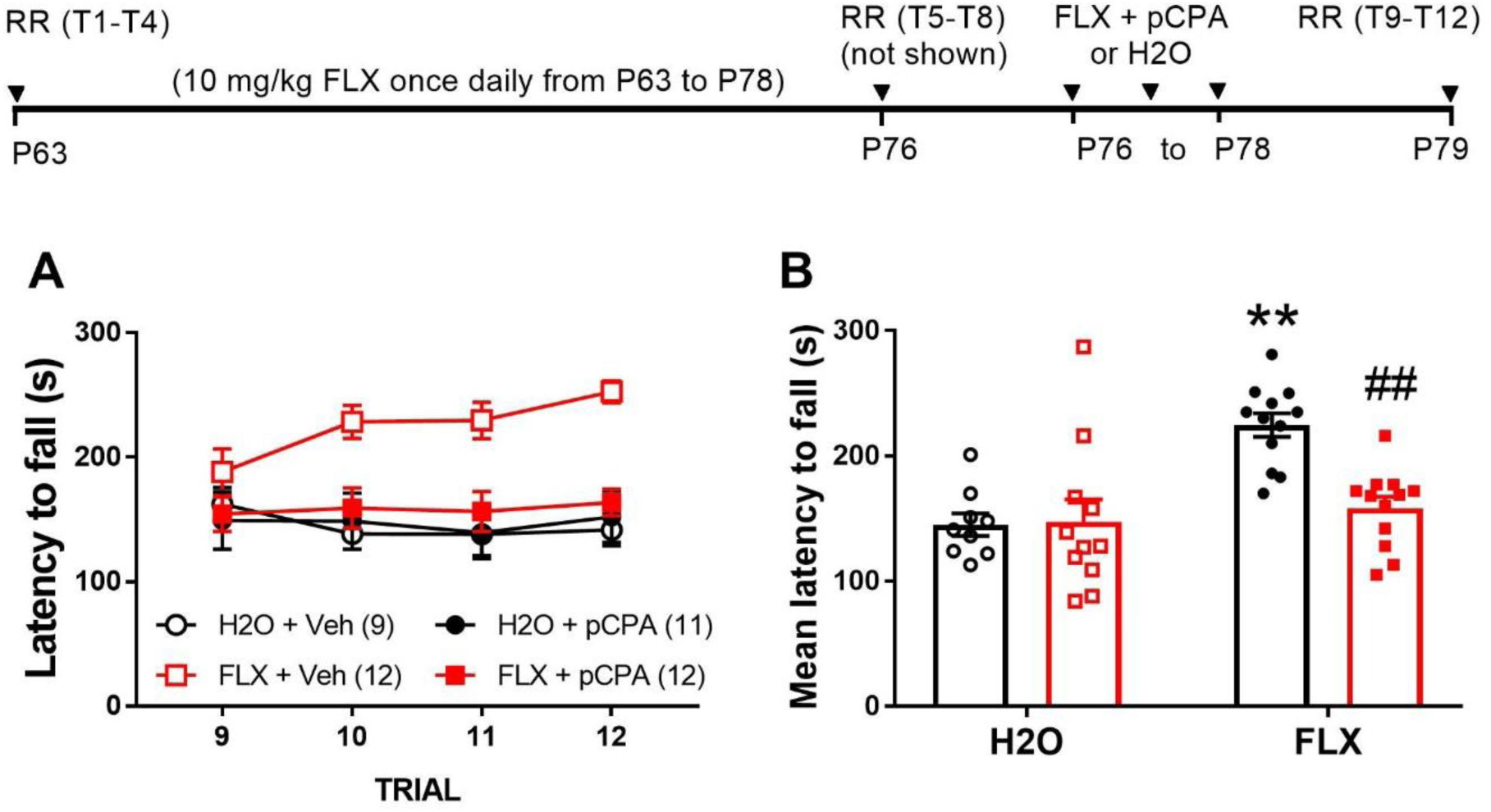
Effect of the 5-HT synthesis inhibitor pCPA on fluoxetine-induced improvement of rotarod performance in Mecp2 Het mice. Panel A show the single trial performance after the administration of fluoxetine alone or fluoxetine + pCPA in *Mecp2* Het mice given repeated dosing of fluoxetine or water. The mean of the 4 daily trials are shown on the right panels. The horizontal bar on top of the figure illustrate the schedule of treatment and timing of behavioral assessment. P, postnatal day; RR, rotarod. Black and red symbols in panel B represent individual data points. Symbols in panel A and histograms in panel B are the mean ± SEM. The number of mice per group is shown in parentheses. **p = 0.0003 vs. H_2_O-Veh, ##p = 0.0011 vs. FLX-Veh (Sidak’s test).

### 3.4 Effects of fluoxetine on rotarod deficit in male Mecp2 null mice

*Mecp2* null mice show severe rotarod deficits already at P35 causing about 50% reduction in the latency to fall as compared to WT mice (Fig. 6). Daily injections of 10 mg/kg fluoxetine for two weeks slightly but significantly increased the latency to fall in *Mecp2* null mice as compared to those receiving vehicle.

**Fig. 6.**
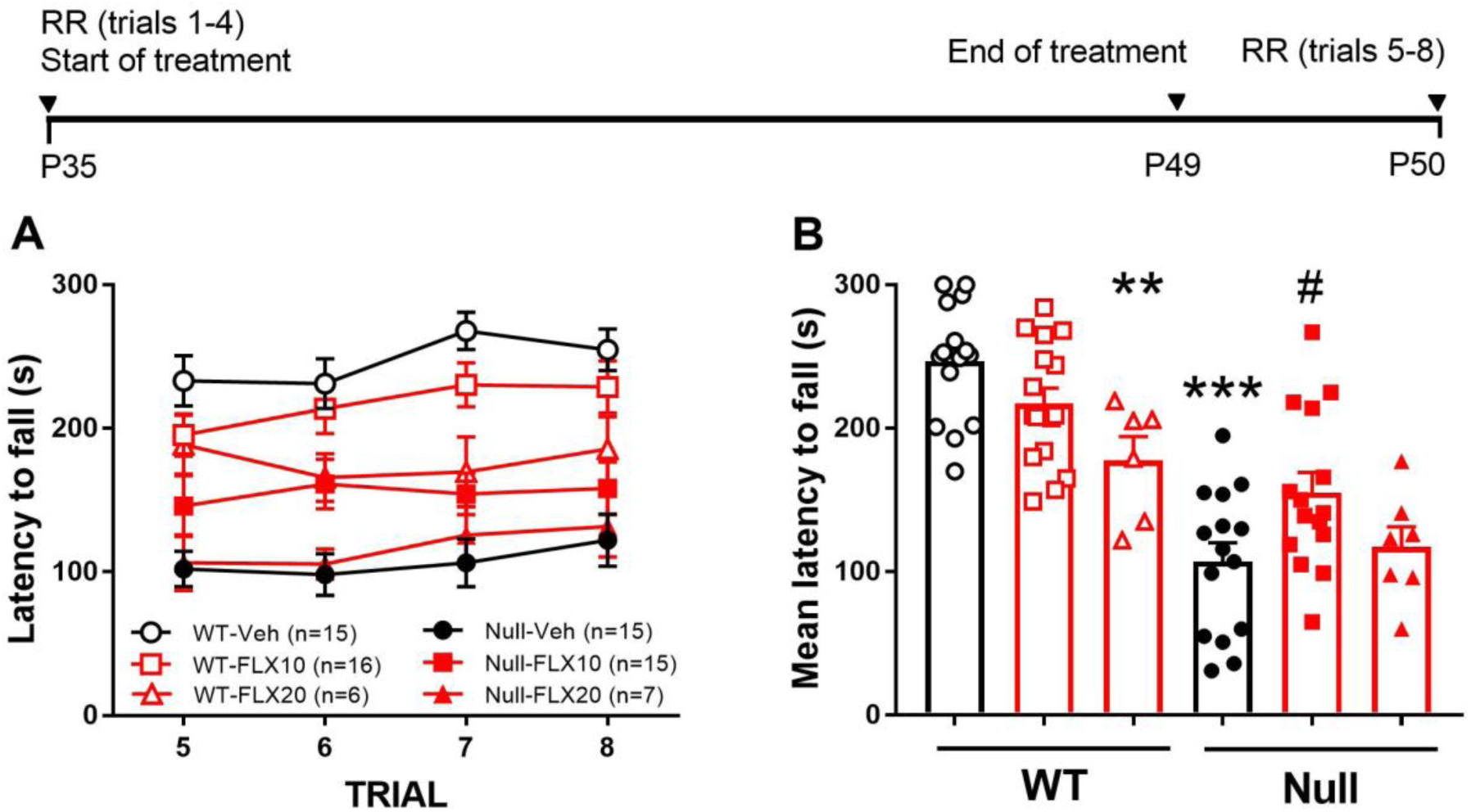
Rotarod performance in Mecp2 null (Null) and wild type (WT) mice treated with 10 and 20 mg/kg fluoxetine (FLX) or vehicle (Veh) for 14 days. The latency to fall for each trial and the mean latency to fall of 4 daily trials are shown in panels A and B, respectively. The horizontal bar on top of the figure illustrate the schedule of treatment and timing of behavioral assessment. Symbols and histograms in panel B represent individual data points and the mean of the group, respectively. Symbols in panel A are the mean ± SEM. The number of mice per group is shown in parentheses. P, postnatal day; RR, rotarod. **p = 0.0082, ***p < 0.0001 vs. WT-Veh; #p = 0.018 vs. Null-Veh (Sidak’s test).

Increasing the dose of fluoxetine to 20 mg/kg had no effect in *Mecp2* null mice, but significantly reduced rotarod performance in WT mice. ANOVA showed a significant effect of treatment (F(2,68) = 3.227, p = 0.0458), genotype (F(1,68) = 56.59, p< 0.0001) and interaction between genotype and treatment (F(2,68) = 6.372, p = 0.0029).

### 3.5 5-HT synthesis

Table1 shows the accumulation of 5-HTP after aromatic aminoacid decarboxylase inhibition, an in vivo indicator of 5-HT synthesis, in various brain regions of male and female *Mecp2* mutants and WT mice. A significant reduction of 5-HTP levels by 31, 25 and 15 % was found in the PFC, HIP and STR of M*ecp2*-null mice, respectively. No significant changes were observed in the BST and RCx. No significant differences in 5-HTP levels were observed between female *Mecp2* Het and WT mice in any of the brain regions examined.

**Table 1.** Accumulation of 5-HTP in various brain regions of *Mecp2*-null, *Mecp2* Het and respective WT mice.

|  | WT | *Mecp2*-null | WT | *Mecp2*-Het |
| --- | --- | --- | --- | --- |
|  | 5-HTP ng/g tissue (n) |  |  |  |
| Prefrontal cortex | 168 ± 18 (6) | **116 ± 14<sup>a</sup>** (6) | 186 ± 10 (6) | 163 ± 14 (6) |
| Hippocampus | 352 ± 15 (7) | **263 ± 12<sup>b</sup>** (6) | 372 ± 21 (6) | 353 ± 46 (6) |
| Striatum | 283 ± 15 (7) | **240 ± 7<sup>c</sup>** (6) | 207 ± 11 (6) | 233 ± 29 (6) |
| Brainstem | 461 ± 36 (7) | 391 ± 28 (6) | 506 ± 23 (6) | 531 ± 39 (6) |
| Remaining Cortex | 199 ± 16 (7) | 168 ± 7 (6) | 211 ± 8 (6) | 199 ± 9 (6) |

Data are mean ± SEM. Significant differences are indicated in bold-type. <sup>a</sup>p = 0.0415, <sup>b</sup>p = 0.0008; <sup>c</sup>p = 0.0309 (Student's t-test)

### 3.6 Effect of fluoxetine on extracellular 5-HT in the PFC

Fig. 7 shows basal extracellular 5-HT in the PFC of WT (panel A) and *Mecp2* mutant (panel B) treated with vehicle or fluoxetine for 6 days. Twenty-four h after the last dose extracellular 5-HT was significantly higher in WT mice receiving fluoxetine as compared to those treated with vehicle. No differences in 5-HT levels were observed between sexes either in mice given vehicle or fluoxetine. Likewise, no differences in extracellular 5-HT were found between *Mecp2* Het and *Mecp2* null mice receiving repeated doses of vehicle. ANOVA showed a highly significant effect of treatment (F(1,14) = 82.97; p < 0.0001) but not sex (F(1,14) = 0.0223; p = 0.883) and interaction between treatment and sex (F(1,14) = 0.185; p = 0.674). Repeated treatment with fluoxetine significantly increased extracellular 5-HT in *Mecp2* Het mice, reaching levels similar to those observed in WT mice receiving the same treatment, whereas fluoxetine had much less effect in *Mecp2* null mice (panel B). ANOVA showed a significant effect of treatment (F(1,18) = 22.21; p = 0.0002) and sex (F(1,18) = 6.412; p = 0.0209) but not interaction between these factors (F(1,18) = 2.915; p = 0.105).

**Fig. 7.**
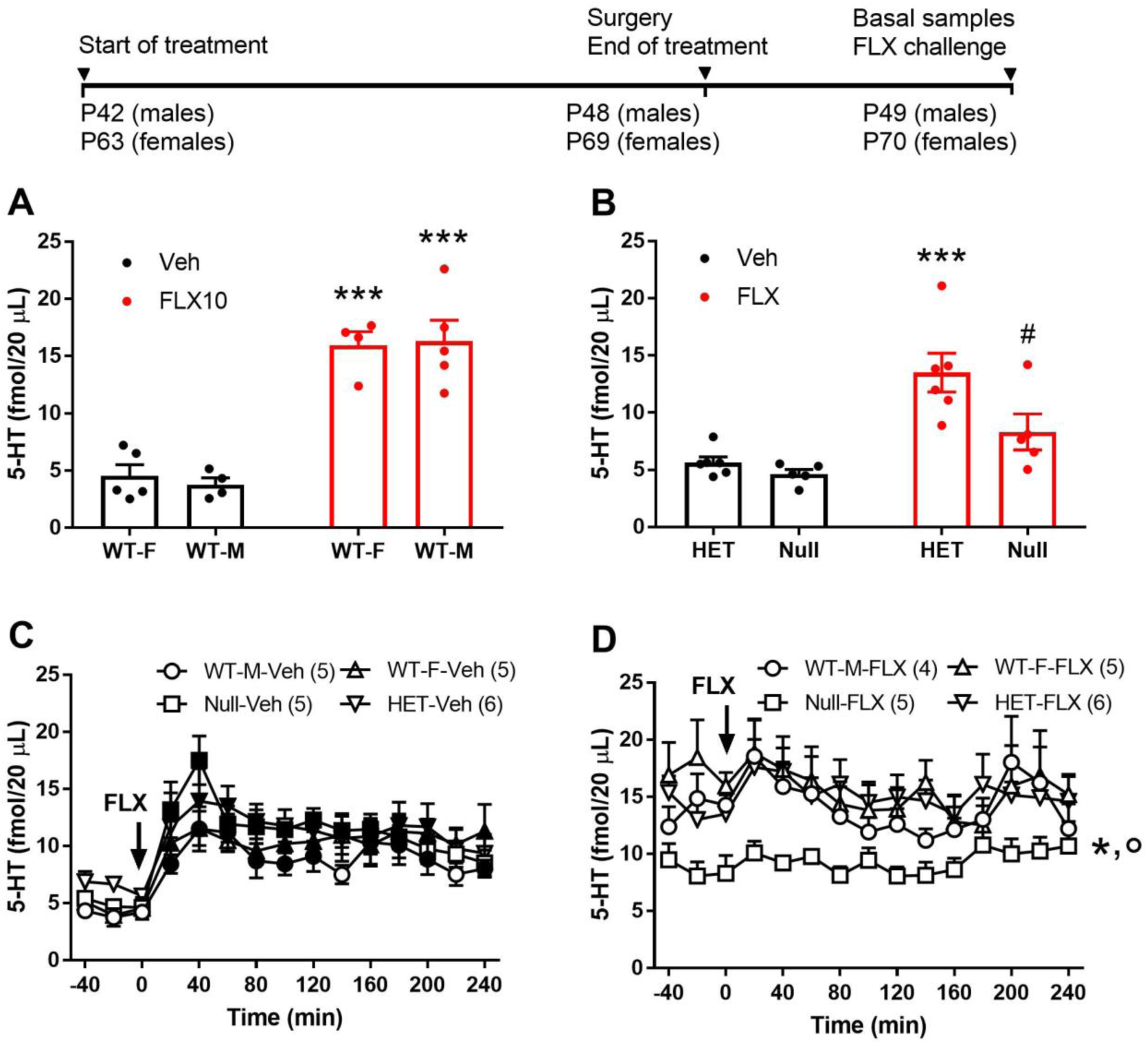
Effect of repeated treatment with fluoxetine on extracellular 5-HT in the PFC. Basal extracellular levels of 5-HT in the PFC of WT male (WT-M) and female (WT-F) mice (panel A), and *Mecp2* Het (HET) female and *Mecp2*-null (Null) male mice (panel B). 5-HT was measured 24 h after 6 days treatment with vehicle (Veh) or fluoxetine (FLX). Dots in panels A and B represent individual data points. ***p < 0.0001 vs. respective vehicle; #p = 0.0032 vs. Het-FLX (Sidak’s test). Lower panels show he effects of a challenge dose of fluoxetine administered 24 h after the last dose of vehicle (Veh; panel C) or fluoxetine (FLX; panel D) in WT and *Mecp2*-mutant mice of both sexes. Histograms in panels A and B and symbols in panels C and D are mean ± SEM. The number of mice/group is shown in parentheses. Solid symbols in panel C indicate p < 0.05 vs. pre-injection (basal) values. *p < 0.05 between Null-FLX and WT-M-H_2_O; °p < 0.05 between Null-FLX and HET-FLX (Tukey’s test).

After an acute fluoxetine challenge, mice repeatedly treated with vehicle show a significant increase of extracellular 5-HT, regardless their genotype and sex (panel C). ANOVA showed significant effects of time (F(1,17) = 17.127; p < 0.0001) but not sex (F1,17) = 0.280; p = 0.604), genotype (F(1,17) = 1.493; p = 0.239) and interaction between genotype and sex (F(1,17) = 0.257; p = 0.619). In mice repeatedly treated with fluoxetine, no further increase was observed upon the administration of the acute challenge in WT and mutant mice of both sexes. Extracellular 5-HT in *Mecp2* null mice remained significantly lower than in WT and *Mecp2* Het mice over the entire observation period (panel D). ANOVA showed significant effects of sex (F(1,15) = 5.023; p < 0.041), time (F(12,180) = 3.703; p < 0.0001), but not genotype (F(1,15) = 2.544; p = 0.132) and interaction between genotype and sex (F(1,15) = 2.186; p = 0.160).

### 3.7 Effects of fluoxetine on 8-OH-DPAT-induced hypothermia

8-OH-DPAT reduced core temperature, reaching −1.7°C in WT male and −1.9°C in WT female mice. 8-OH-DPAT had similar effect in *Mecp2* null mice (−1.6°C), whereas the drop in temperature was significantly greater in *Mecp2* Het mice (−2.6°C; Fig. 8). The hypothermic effect induced by 8-OH-DPAT was markedly attenuated by fluoxetine in male and female mice of both genotypes. In male mice, ANOVA showed significant effects of treatment (F(1,21) = 4.989; p = 0.037) but not genotype (F(1,21) = 0.023; p = 0.882), and interaction between genotype and treatment (F(1,21) = 0.065; p = 0.801). In female mice, ANOVA showed significant effects of genotype (F(1,21) = 7.498; p = 0.0123), treatment (F(1,21) = 61.41; p < 0.0001) but not interaction between genotype and treatment (F(1,21) = 1.043 p = 0.319).

**Fig. 8.**
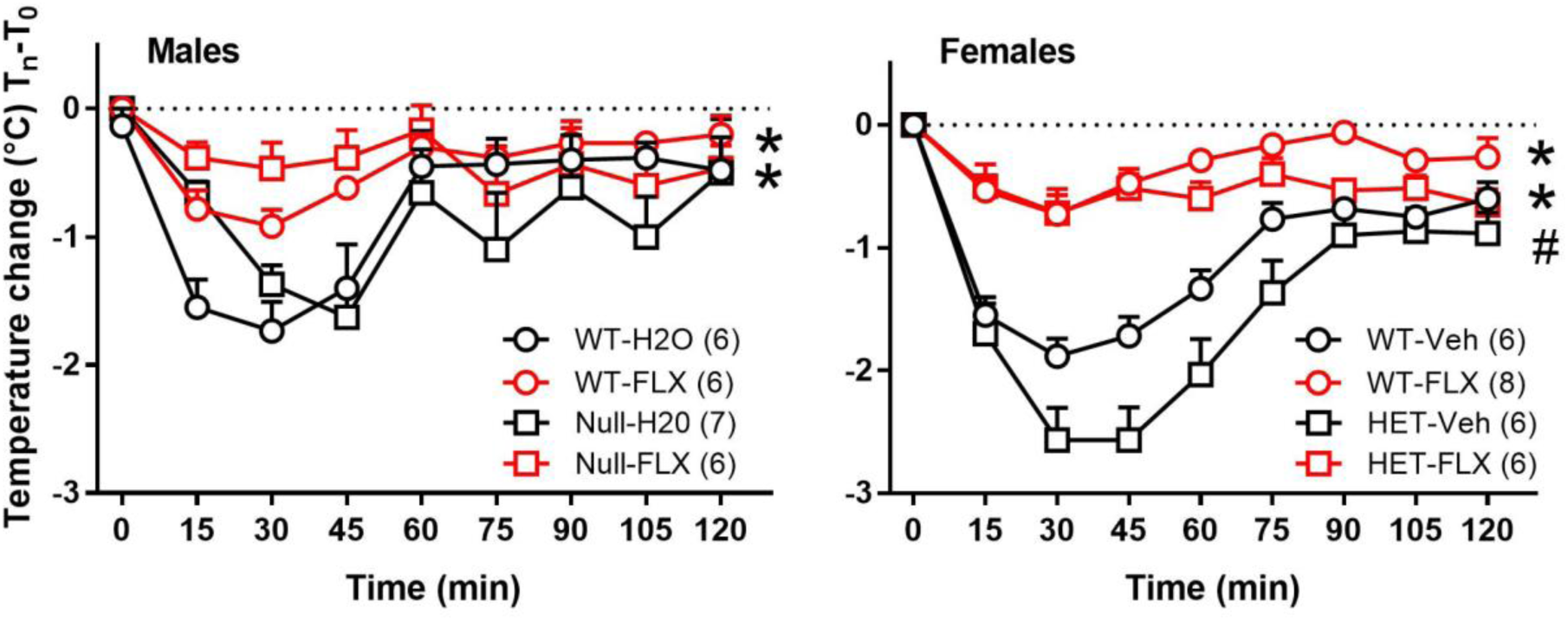
Effect of 6 days treatment with fluoxetine (FLX) or vehicle (Veh) on 8-OH-DPAT-induced hypothermia. Male *Mecp2*-null (Null) and WT mice and female *Mecp2*-Het (HET) and WT mice were injected with 8-OH-DPAT at t = 0 min. Core temperature was measured immediately before 8-OH-DPAT injection and every 15 min thereafter. Data are expressed as mean ±SEM of the difference between temperature measured at each time after 8-OH-DPAT injection and basal values (T_0_). Basal temperature (°C) measured 24 h after the repeated administration of fluoxetine or vehicle for 6 days was: males, WT-Veh 38.2 ± 0.2, WT-FLX 38.3 ± 0.1, *Mecp2* null-Veh 37.7 ± 0.2 and *Mecp2* null-FLX 38.0 ± 0.3; females, WT-Veh 38.2 ± 0.1, WT-FLX 38.1 ± 0.1, *Mecp2* Het-Veh 38.0 ± 0.1 and *Mecp2* Het-FLX 38.3 ± 0.1.

The number of mice/group is shown in parentheses. *p <0.05 between WT-FLX and WT-H_2_O; °p < 0.05 between Het/null-FLX and Het/null-Veh; #p < 0.05 between Het-Veh and WT-Veh (Tukey’s test).

## 4. Discussion

Motor deficits represent one of the most disabling symptoms of RTT. *Mecp2* Het mice show an early onset of rotarod deficit (Samaco et al., 2013), worsening over time, which is fully rescued by fluoxetine treatment. The rotarod improvement was larger than commonly found with other molecules tested so far in mouse models of RTT (Braun et al., 2012; Buchovecky et al., 2013; Krishnan et al., 2015; Mellios et al., 2014; Nag and Berger-Sweeney, 2007) and comparable to that observed after genetic rescue of *Mecp2* gene (Chao et al., 2010; Garg et al., 2013). The robustness of our findings is supported by the ability of fluoxetine to rescue rotarod deficit in *Mecp2* Het mice differing for age, training on the rotarod, severity of motor impairment, and route of drug administration (drinking water; Fig. S2). The ability of fluoxetine to rescue the coordination deficit was confirmed in the beam walking task, that similarly to rotarod assesses motor coordination and balance (Carter et al., 2001). Likewise, citalopram rescued rotarod and beam walking deficits. Although the number of slips of *Mecp2* Het mice used to assess the effect of fluoxetine was larger than in the experiment with citalopram, these differences apparently had no consequences on drugs’ ability to counteract beam walking deficit. Previous studies showed that i.p. injection, the route used for fluoxetine administration, increased corticosterone secretion and influenced neuronal activity and mouse behavior (Cabib et al., 1990; Ryabinin et al., 1999; Sousa et al., 2006; Stuart and Robinson, 2015). Thus, the different number of slips showed by *Mecp2* Het mice across experiments, is likely related to the stress caused by the i.p. injection, which depending on mouse genetic background may adapt or not to repeated injections (Ryabinin et al., 1999). Fluoxetine failed to improve performance in the hanging-wire test. As muscle strength is required to oppose the effect of gravity in the hanging-wire but not in the rotarod and beam walking tests, it is conceivable that fluoxetine mainly affects motor coordination and balance rather than muscle strength (Klein et al., 2012; Osmon et al., 2018). Further assessment using other tests, such as grip strength, are needed to support this interpretation. However, it cannot be excluded that extending treatment duration by more than 2 weeks or starting fluoxetine administration before symptom onset may improve the hanging-wire performance.

We found that 5-HT synthesis inhibition abolished the rotarod improvement induced by fluoxetine in *Mecp2* Het mice indicating a major role of 5-HT in this effect. The fact that citalopram, another SSRI with a different chemical structure, shared the ability of fluoxetine to rescue motor deficits in the rotarod and beam walking tests further supports the importance of 5-HT in the mechanisms by which these drugs improve motor coordination. The dose of pCPA used in the current study reduced 5-HTP accumulation by 50-75% and prevented the antidepressant-like effect of citalopram and paroxetine in the forced swimming test (Cervo et al., 2005; Guzzetti et al., 2008). In addition, polymorphism of the *Tph2* gene, which reduces brain 5-HT synthesis by 25-40%, was sufficient to prevent the antidepressant-like effect of SSRIs (Cervo et al., 2005; Guzzetti et al., 2008). These findings are consistent with previous studies showing that SSRIs enhance motor skill learning and plasticity likely by enhancing synaptic availability of brain 5-HT, which is known to regulate motor networks and facilitate motor skill (Batsikadze et al., 2013; Gerdelat-Mas et al., 2005; Loubinoux et al., 2005).

We found that fluoxetine was poorly effective in contrasting rotarod deficits in male *Mecp2* null mice. Interestingly, we found reduced 5-HT synthesis in the PFC, HIP and STR of male *Mecp2*-null mice, while no changes are found in no brain region of *Mecp2* Het mice. These findings are consistent with data showing reduced 5-HT levels, and *Tph2* gene expression and protein in the raphe and other brain regions of *Mecp2* null mice and in post-mortem brain samples of RTT patients (Ide et al., 2005; Panayotis et al., 2011; Samaco et al., 2009; Santos et al., 2010; Vogelgesang et al., 2017). Except for a study showing reduced levels of 5-HT in the raphe region but not in the hippocampus (Isoda et al., 2010), evidence for brain 5-HT alterations in *Mecp2*-deficient female mice are lacking. In all the brain areas examined for changes in 5-HTP accumulation, tryptophan levels were similar between *Mecp2* null and WT mice (Table S1). Thus, we can exclude that insufficient supply of 5-HT precursor TRP from diet may account for reduced 5-HT synthesis (Fernstrom, 1983). Neurodegeneration of 5-HT neurons is also excluded as no differences in the number of brain 5-HT neurons have been observed between *Mecp2*-null and WT mice (Santos et al., 2010). Thus, the most parsimonious explanation for reduced 5-HT synthesis in *Mecp2* null mice is a cell autonomous mechanism exerted through the direct control of *Tph2* expression by the *Mecp2* gene Samaco et al., 2009).

Differences in brain and plasma levels of fluoxetine, as well as differences in norfluoxetine to fluoxetine ratio have been found between male and female mice (Hodes et al., 2010). In agreement, we found lower brain and plasma levels of nor-fluoxetine in *Mecp2*-null mice as compared to mutant females (Table S2). However, doubling the dose of fluoxetine failed to improve rotarod performance in *Mecp2*-null mice despite plasma and brain fluoxetine and norfluoxetine reached levels similar or higher than those found in females receiving 10 mg/kg fluoxetine (Table S2). Thus, the poor effect of fluoxetine in male *Mecp2*-null mice is not attributable to low exposure to the drug or its metabolite. It is unlikely that increasing further fluoxetine dose would improve its efficacy in male mice as reduced rotarod performance is observed in mice receiving 20 mg/kg fluoxetine (present study; Fernández-Guasti et al., 2017; Morelli et al., 2011).

The motor effect of fluoxetine was no more present 3 days after drug suspension. This indicates that sufficient amounts of fluoxetine and its active metabolite should be present in the brain to support the effect of fluoxetine. The half-life of fluoxetine and nor-fluoxetine in the mouse plasma (about 6 and 12 h, respectively) and the time-dependent binding of the drug to the 5-HT transporter are consistent with this interpretation (Hirano et al., 2005).

Fluoxetine administration for at least 6 days is required to achieve a significant rescue of rotarod deficit. The effect is maintained during 2 weeks of administration but is not achieved by a single dose or administration for 3 consecutive days. Likewise, 1-week administration of fluoxetine, a condition sufficient to achieve steady-state levels of brain and circulating fluoxetine (Miller et al., 2008), enhanced extracellular 5-HT in the PFC of *Mecp2* Het significantly more than in *Mecp2*-null mice. On the contrary, neither basal 5-HT release nor the effect of a single dose fluoxetine on extracellular 5-HT is affected by sex and genotype. Consistently, SERT expression in the PFC, motor cortex and brainstem is not affected by *Mecp2* gene deletion (Santos et al., 2009; Vogelgesang et al., 2017). However, reduced expression of SERT was found in the hippocampus (Vogelgesang et al., 2017). Thus, it cannot be excluded that basal and fluoxetine-induced enhancement of extracellular 5-HT may be affected in extracortical regions of *Mecp2* mutant mice. The development of adaptive mechanisms in response to prolonged fluoxetine administration is responsible for the delayed effect of SSRIs in enhancing the availability of extracellular 5-HT in the PFC and striatum. This occurs mainly through the down-regulation of 5-HT_1A_ autoreceptors-mediated autoinhibitory feedback (Chaput et al., 1991; Fritze et al., 2017; Hensler, 2002; Invernizzi et al., 1996). These prompted us to assess whether sex-dependent adaptation of 5-HT_1A_ autoreceptors sensitivity may account for the inability of fluoxetine to restore rotarod performance in *Mecp2* null mice. To address this point, we assessed the hypothermic effect of 8-OH-DPAT, a red-out of 5-HT_1A_ autoreceptors sensitivity (Goodwin et al., 1985). We found that fluoxetine desensitized 5-HT_1A_ autoreceptors to a similar extent in *Mecp2* null and *Mecp2* Het mice. Thus, the reduced efficacy of fluoxetine on motor coordination and extracellular 5-HT in *Mecp2* null mice is not attributable to sexually dimorphic changes in 5-HT_1A_ autoreceptors sensitivity. However, we found that the hypothermic effect of 8-OH-DPAT was potentiated in *Mecp2* Het, but not in *Mecp2* null mice. Enhanced sensitivity to the hypothermic effect of 8-OH-DPAT were also observed in male mice selectively lacking the *Mecp2* gene in brain serotonergic neurons (Philippe et al., 2018). These mice showed increased expression, binding and immunoreactivity of 5-HT_1A_ autoreceptors in the raphe nuclei and sex-dependent alteration of emotional behavior (Philippe et al., 2018). These findings suggest that increased expression of 5-HT_1A_ autoreceptors may also underlie the enhanced sensitivity to the hypothermic effect of 8-OH-DPAT in *Mecp2* Het mice. Increased expression and sensitivity of 5-HT_1A_ autoreceptors, may result in autonomic dysregulation of respiratory and cardiac rhythms (Audero et al., 2008; Corcoran et al., 2013), which are prominent and life-threatening symptoms affecting nearly 100% of subject with RTT over the lifespan (Tarquinio et al., 2018). Thus, fluoxetine may help counteract respiratory deficits of RTT by desensitizing 5-HT_1A_ autoreceptors. Consistently, breathing irregularities in a girl with RTT were reduced by fluoxetine (Gokben et al., 2012).

### Conclusions

The present study provides robust evidences showing that fluoxetine and citalopram rescue motor coordination deficits in *Mecp2* Het mice and that brain 5-HT is necessary for these effects. These findings suggest that fluoxetine and potentially other SSRIs and drugs enhancing 5-HT neurotransmission may have beneficial effects on motor and potentially other symptoms of RTT. However, the pleiotropic effects of fluoxetine encompassing 5-HT neurotransmission, neurogenesis, synaptic plasticity, expression of neurotrophic factors and facilitation of MeCP2 protein expression, which are all defective in RTT, suggest that in addition to 5-HT other factors may contribute to the effect of fluoxetine on motor coordination (Cassel et al., 2006; Chang et al., 2006; Duman and Monteggia, 2006; Maya Vetencourt et al., 2008, Nag et al., 2013; Santarelli et al., 2003; Stagni et al., 2015; Villani et al., 2018). Further studies in animal models more closely reflecting the molecular alterations of the *Mecp2* gene found in patients with RTT are needed to improve the translational validity of the present findings and provide the basis to assess whether fluoxetine may have any place in the therapy of RTT.

## Acknowledgments

We thank Dr P. Gagliardi for helping with behavioral and biochemical experiments

## Supplementary Tables and Figures

**Supplementary Table S1.** Tryptophan levels in various brain regions of male *Mecp2*-null and WT mice.

|  | WT | *Mecp2*-null |
| --- | --- | --- |
|  | Tryptophan ng/g tissue (n) |  |
| Prefrontal cortex | 4677 ± 1038 (6) | 4413 ± 484 (6) |
| Hippocampus | 4396 ± 670 (7) | 4658 ± 788 (6) |
| Striatum | 5207 ± 834 (7) | 5793 ± 961 (6) |
| Brainstem | 4047 ± 755 (7) | 4185 ± 698 (6) |
| Remaining Cortex | 5093 ± 830 (7) | 5357 ± 1030 (6) |

Data are mean ± SEM.

**Supplementary Table S2.** Fluoxetine (FLX) and norfluoxetine (NFLX) levels in plasma and brain of *Mecp2*-deficient mice

| Genotype | FLX dose<br>(mg/kg) | PLASMA |  | BRAIN |  |
| --- | --- | --- | --- | --- | --- |
|  |  | FLX<br>(ng/mL) | NFLX | FLX<br>(µg/g) | NFLX |
| *Mecp2* Het | 10 | 1329±97 | 1690±147 | 17.1±1.6 | 21.7±2.0 |
| *Mecp2* null | 10 | 1076±132 | 1053±35<sup>*</sup> | 15.1±1.0 | 13.8±1.6<sup>**</sup> |
| *Mecp2* null | 20 | 1885±246 | 1636±142 | 37.0±2.2<sup>***</sup> | 31.9±1.5<sup>**</sup> |

Plasma and brain levels of fluoxetine and norfluoxetine were measured in *Mecp2* Het and *Mecp2* null mice receiving 10 mg/kg fluoxetine for 7 days. Mice were sacrificed 4 h after the last dose of the repeated schedule. Plasma in *Mecp2* null mice given 20 mg/kg FLX, was obtained from blood withdrawn from the maxillary sinus after 7 days of repeated injections. Brain tissue was collected from the same mice after 2-week treatment, 4 h after the last dose of FLX. Data are mean ± SEM of 6-7 mice per group. \*p < 0.05, \*\*p < 0.005, \*\*\*p < 0.0001 vs. *Mecp2* Het (Dunnett's test).

**Fig. S1A.**
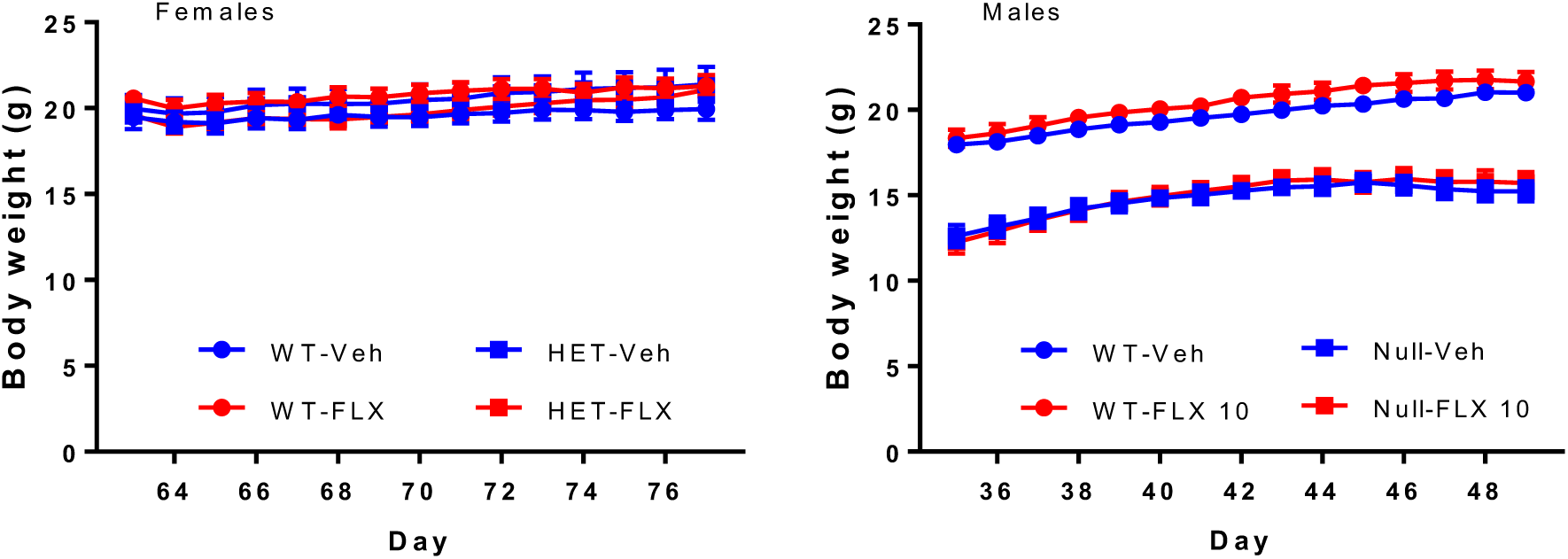
Body weight of *Mecp2* HET and WT female and *Mecp2* null and WT male mice over the 14-day treatment with 10 mg/kg fluoxetine (FLX) or vehicle (Veh), once daily. The same mice were evaluated in the rotarod as reported in Figs. 2 and 6, respectively.

**Fig. S1B.**
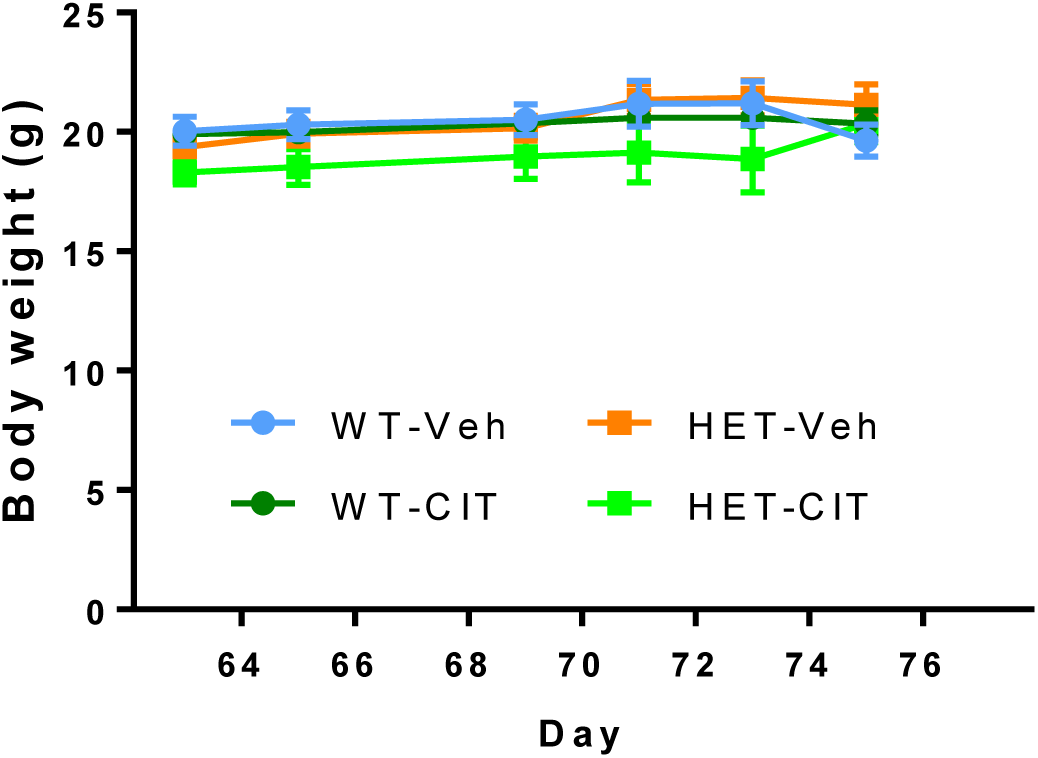
Body weight of *Mecp2* Het and WT mice receiving 20 mg/kg citalopram in drinking water or plain water. The same mice were evaluated in the rotarod as reported in in Fig. 2

**Fig. S2.**
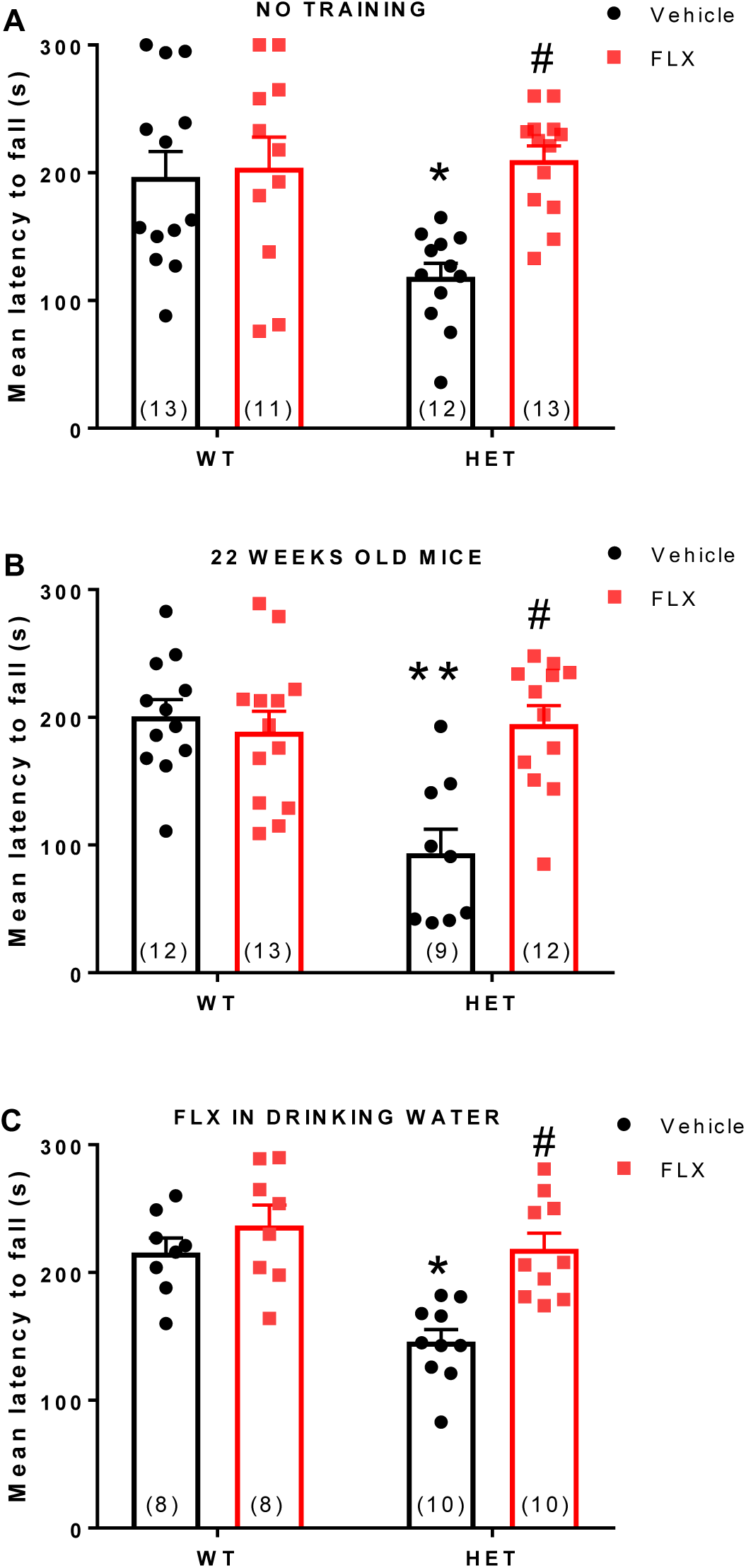
The ability of fluoxetine in reversing motor deficits of *Mecp2* Het (HET) mice is maintained under different experimental conditions: A) mice not pre-trained on the rotarod (ANOVA for treatment F(1,45) = 8.427, p = 0.0057; genotype F(1,45) = 4.532, p = 0.0388; genotype x treatment F(1,45) = 6.137, p = 0.0171); B) 22 weeks old mice (ANOVA for treatment, F(1,42) = 8.055, p = 0.0070; genotype F(1,42) = 10.4, p = 0.0024; genotype x treatment F(1,42) = 12.92, p = 0.0008); C) after administration of FLX in drinking water treatment (ANOVA for treatment, F(1,32) = 14.3, p = 0.0006; genotype F(1,32) = 12.6, p = 0.0012; genotype x treatment F(1,32) = 4.321, p = 0.0457). WT and HET mice were given 10 mg/kg/day fluoxetine or vehicle for 14 days. Data are the mean ± SEM. The number of mice per group is shown in parentheses. *p = 0.002, **p < 0.0007 vs. WT-Veh; #p < 0.02 vs. HET-Veh (Sidak’s test).

